# DVPNet: A New XAI-Based Interpretable Genetic Profiling Framework Using Nucleotide Transformer and Probabilistic Circuits

**DOI:** 10.64898/2026.01.28.695053

**Authors:** Taishi Kusumoto

## Abstract

In this study, we present an XAI-based genetic profiling framework that quantifies gene importance for distinguishing cancer cells from normal cells based on an interpretable AI decision process. We propose a new explainable AI (XAI) classification model that combines probabilistic circuits with the Nucleotide Transformer. By leveraging the strong feature-extraction capability of the Nucleotide Transformer, we design a tractable classification framework based on probabilistic circuits while preserving probabilistic interpretability.

To demonstrate the capability of this framework, we used the GSE131907 single-cell lung cancer atlas and constructed a dataset consisting of cancer-cell and normal-cell classes. From each sample, 900 gene types were randomly selected and converted into embedding vectors using the Nucleotide Transformer, after which the classification model was trained. We then extracted class-specific probabilistic contributions from the tractable model and defined a contribution score for the cancer-cell class. Genetic profiling was performed based on these scores, providing insights into which genes and biological pathways are most important for the classification task. Notably, 1,524 of the 9,540 observed genes showed contribution scores that contradicted what would be expected from their class-wise occurrence frequencies, suggesting that the profiling goes beyond simple statistics by leveraging biological feature representations encoded by the Nucleotide Transformer. The top-ranked genes among these contradictory cases include several well-studied genes in cancer research (e.g., ITGA5, SIGLEC9, NOTUM, and TP73). Overall, these analyses go beyond traditional statistical or gene-expression-level approaches and provide new academic insights for genetic research.

## 1 Introduction

Gene co-expression networks constructed from RNA sequencing data have been used to infer gene functions and disease associations.[1] Indeed, previous studies have identified disease-associated gene modules and biologically meaningful pathways using co-expression network analysis, including in cancer [2] [3], neurological disorders [4], and complex traits.[5] However, gene co-expression networks, constructed from correlational relationships among RNA expression levels, primarily indicate which genes are active in the same biological processes but do not normally provide information about causality or distinguish between regulatory and regulated genes.[1] For instance, genes in the same biological pathway often do not show similar RNA expression patterns, which cannot be adequately treated in co-expression networks. [6] The fundamental reliance of co-expression networks on statistical correlations of expression levels restricts their ability to capture functional, regulatory, or context-dependent relationships among genes. Therefore, due to these limitations, further advancement of genetic research requires a new workflow to develop alternative genetic networks that provide different biological insights beyond statistical analyses of RNA expression levels.

The recent advancement of large language models clearly demonstrates that training transformer-based models on large amounts of text in a self-supervised manner enables strong contextual understanding of arbitrary input sequences, allowing models to solve diverse advanced tasks, including mathematical reasoning, scientific discussions, translation, and coding. [7] Similarly, a foundation model such as the Nucleotide Transformer [8], trained on large collections of nucleotide sequences for next-base prediction in a self-supervised manner, is expected to acquire contextual understanding of biological functions encoded in nucleotide sequences. Such genetic foundation models can encode expressed genes in RNA sequence data into meaningful feature representations that capture inherent biological functions. Indeed, previous studies have shown their usefulness as encoders for downstream tasks.[8] Therefore, utilizing these representations can contribute to the construction of new types of genetic networks that provide biological insights distinct from those obtained through co-expression networks.

In this paper, we adopt the Nucleotide Transformer as an encoder for genes in RNA sequence data. We reconstruct nucleotide sequences encoding each gene by working backwards from transcription start sites (TSS). These nucleotide sequences are then input to obtain embedded feature representations for each expressed gene in a cell sample. Directly encoding mature RNA sequence data may lose information related to intronic regions; however, this approach avoids such information loss. The resulting embedded vectors are integrated into a novel explainable gene classification model, DVPNet, which is derived from VPNet, an explainable image classification model that combines embedded vectors from vision transformers with probabilistic circuits.

Gene classification models [9] can extract and select important features and internally identify genes that are important for specific classification tasks, such as distinguishing cancer cells from normal cells. However, the black-box nature of traditional classification models based on intractable neural networks, including convolutional neural networks [10] and transformers [11], complicates the interpretation of their decision processes. In contrast, DVPNet is designed to encode conditional probabilistic contributions for each gene to each class, given a sample, while leveraging the biological information encoded by the Nucleotide Transformer. These probabilistic contributions can be utilized for downstream analyses, including the construction of new genetic networks, enabling biological analyses from perspectives different from those provided by co-expression networks. In this paper, we demonstrate the construction of gene-level contribution landscapes and show that downstream biological analyses can be performed, supporting the validity of this workflow. This workflow can be applied to a wide range of biological classification tasks and has the potential to complement existing workflows in genetic research.

## 2 Methods

### 2.1 VPNet

VPNet [12] combines a tractable probabilistic circuit [13] with patch-embedding vectors from vision transformers.[14][15][16] The study demonstrated that, by leveraging the strong feature-extraction capability of ViTs, the model achieves significant robustness and representational capacity without losing tractability, by structurally preserving smoothness and decomposability.

VPNet is a novel image-classification algorithm that can probabilistically visualize the decision process of the classification model. Unlike most traditional classification models that adopt convolutional neural networks (CNNs) or transformers, the algorithm combines a vision transformer with probabilistic circuits, preserving tractability and explainability. This model can encode distributions over several independent sources of information. For instance, when the model treats images with two modalities, two encoders, and two classes, each probabilistic circuit in the model encodes the following class-conditional densities:

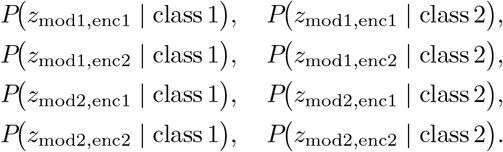

Here, each class-conditional density can be combined (under the model’s factorization) into the joint class-conditional density over all information:

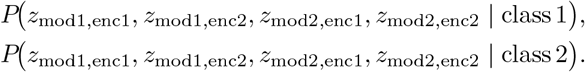

This means that VPNet can be optimized at the per-sample level, while at inference it can extract any partial information included in the sample. This enables visualization of the probabilistic contributions for each piece of information in the classification task. Figure 1 is one example of such a visualization. In the figure, the heatmaps clearly highlight the defective regions for 2D image classification between defective and non-defective products, demonstrating the validity of the VPNet algorithm. The more detailed description of the algorithm is described in [12].

**Figure 1.**
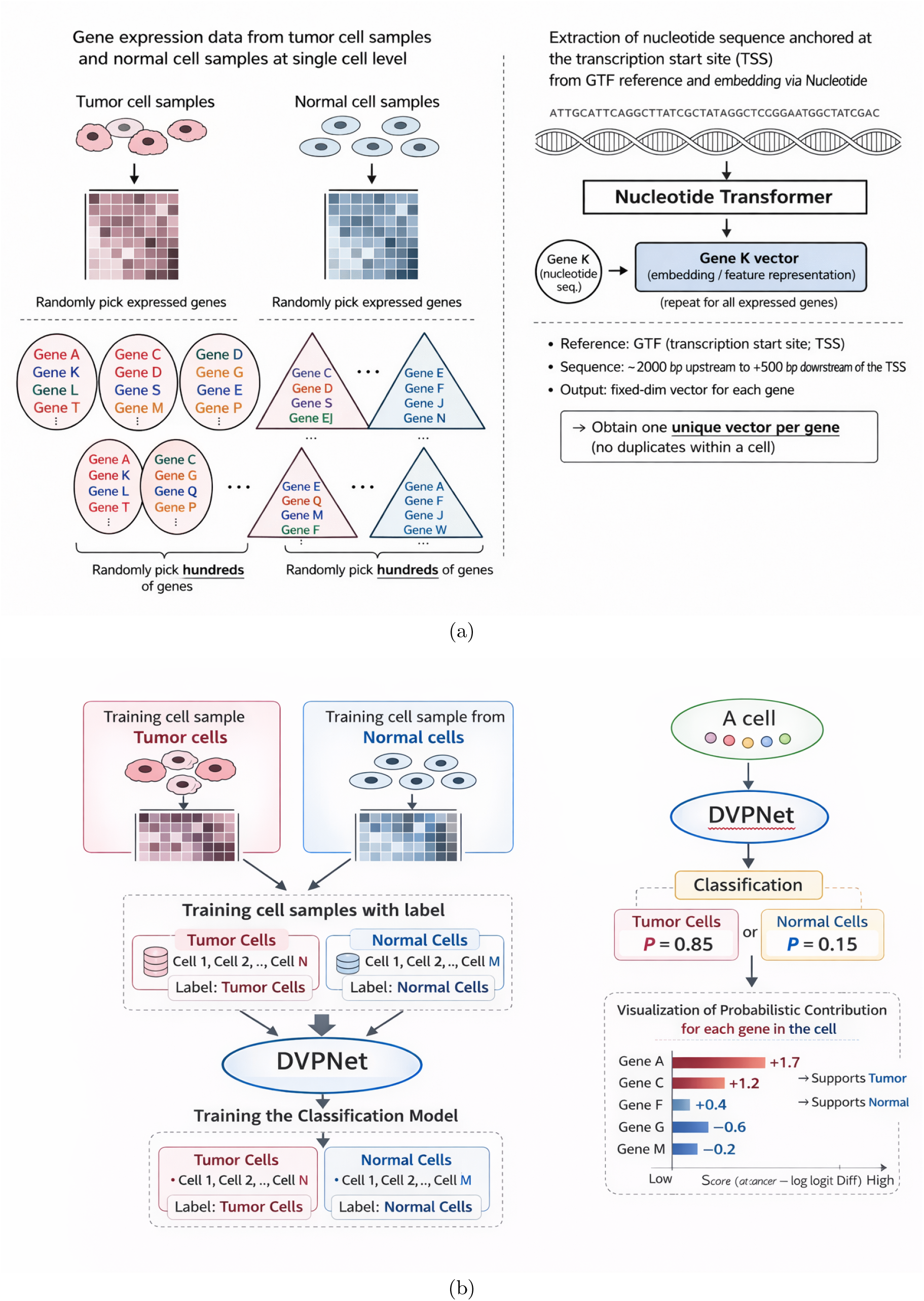
Overall experimental workflow: (a) Sample preparation using the Nucleotide Transformer, and (b) model training and extraction of gene-wise probabilistic contributions for each sample.

### 2.2 Encoding gene-level vectors using Nucleotide Transformer

Nucleotide Transformer is a foundation model optimized for the masked genetic-sequence prediction task. Like other large models (LMs), including large language models and vision transformers, Nucleotide Transformer is trained on a large amount of nucleotide sequences from 3,202 human genomes and 850 genomes from diverse species in an unsupervised manner. This provides a guarantee of robust representational learning for structures, functions, and relationships. Therefore, this foundation model can be used to extract feature representations for downstream tasks.

From a single-cell–level RNA expression dataset, we extract *N* RNAs without any name overlap in each single cell. Then, for each RNA, the corresponding transcription start site (TSS) is determined using the reference genome annotation in Gene Transfer Format (GTF), and a nucleotide sequence of fixed length anchored at the TSS is extracted. In this paper, we extracted the nucleotide sequences from -2000 bp upstream to +500 bp downstream of the TSS. The extracted sequence is input to Nucleotide Transformer, and the embedding matrix containing each token vector from the second-to-last layer is obtained. Finally, the embedding matrix is converted to an embedding vector by average pooling of each scaler component of all token vectors. We call this vector a *gene vector*. The gene vector replaces the patch-embedding vector in VPNet; equivalently, we replace the encoder (vision transformer) in VPNet with Nucleotide Transformer.

### 2.3 Single cell level gene extraction

We assumed that gene regulation can be influenced by genes with low RNA expression levels as well as by genes with high RNA expression levels. Therefore, all expressed genes in a single cell were initially considered in this study, regardless of their expression levels. However, simply using all expressed genes in a cell may cause the model to focus primarily on statistical differences in the frequencies of gene types across classes. Because the model should capture intrinsic functional signals beyond such statistics, while still leveraging the meaningful embeddings produced by the Nucleotide Transformer, we randomly sampled a subset of 900 expressed genes per cell. This uniform random sampling treats genes with low RNA expression levels and genes with high RNA expression levels equally. Figure 1 summarizes the workflow.

### 2.4 DVPNet: gene-level explainable classification algorithm

By replacing the vision transformer with the Nucleotide Transformer, VPNet is remodeled into a gene-level classification algorithm, which we call *DVPNet*. Each sample comprises 900 gene vectors with no overlap:

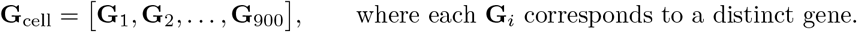

As we pick up these genes without overlap, this ensures that gene-vector scopes are disjoint, satisfying decomposability:

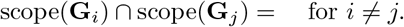

Each embedding vector **G** is an input unit to the probabilistic circuit. Each **G** comprises 1024 scalar components,

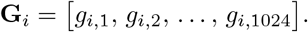

Additionally, during optimization, the parameters of the Nucleotide Transformer are frozen, so the scope of each scalar component is invariant. In particular,

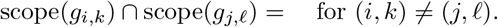

This construction ensures decomposability for product nodes, and smoothness is enforced by defining sum-node children to have identical scopes.

Then, each scalar component of a vector is input to a leaf node of the probabilistic circuit, as in VPNet. The circuit prepares one leaf node for each scalar component. This input is encoded as a univariate probability density, such as a Gaussian,

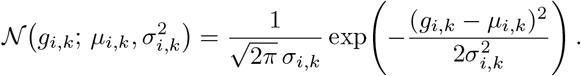

As in VPNet, smoothness and decomposability are strictly preserved in the structure of our probabilistic circuit. Consequently, the circuit encodes the joint probability distribution over all scalar components; because the Nucleotide Transformer is frozen during optimization, each gene vector has an immutable scope inherited from its corresponding gene. Thus,

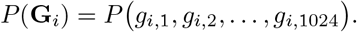

Then, we factorize the joint probability over all gene vectors included in the sample. Because gene-vector scopes are disjoint, this preserves decomposability, which means that the result of the factorization equals the joint probability of all genes in the sample. Additionally, the joint probability distribution over all genes equals the joint probability over the sample scope:

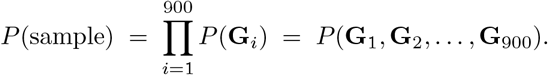

In VPNet, the number of prepared probabilistic circuits is defined as

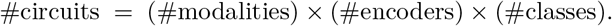

Here, we use one modality (genes), one encoder (Nucleotide Transformer), and two classes in all experiments. Thus, the number of probabilistic circuits per experiment is 2. We treat the output of each probabilistic circuit as a probabilistic model encoding the *class-conditional* probability of the sample (class 1):

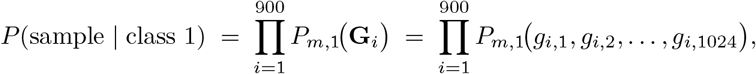

where *P*_*m*,1_ denotes the circuit (for modality *m* and class 1) that factorizes over the scalar components of each gene vector.

Similarly, the output of the other circuit is treated as the class-conditional probability for class 2:

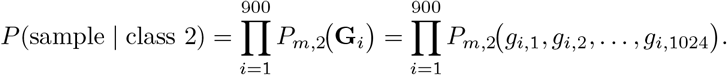

As in VPNet, we parameterize the class prior with a small neural network as a Bernoulli distribution:

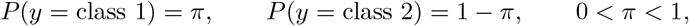

where*π* is a learnable parameter.

Then, by Bayes’ rule,

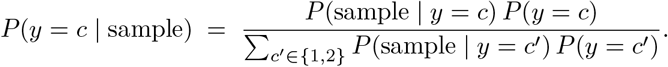

However, to stabilize the training process, we adopted the geometric mean of the likelihoods across the 900 feature vectors instead of the raw product likelihood. We define

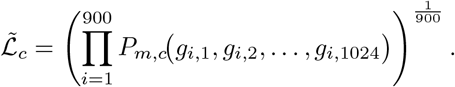

Then, a power posterior is computed by applying Bayes’ rule to the geometric mean likelihood:

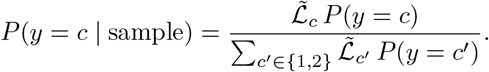

Although this departs from the exact Bayesian posterior under the full product likelihood, it preserves a valid probabilistic interpretation as a normalized power posterior [17].

As in VPNet, optimization is performed per sample. Let

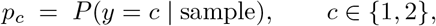

and let **y** = (*y*_1_, *y*_2_) be the one-hot target. The loss combines cross-entropy with a Shannon-entropy regularizer:

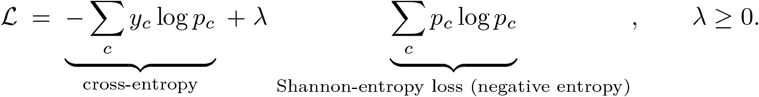

In the training dataset, some samples may share the same gene (i.e., the same scope), but no product node connects any samples; thus, decomposability of the model is preserved during optimization. Additionally, two gene vectors from the same gene can differ because standardization is computed over the vectors in each *sample*:

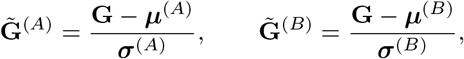

where (***µ***^(*A*)^, ***σ***^(*A*)^) and (***µ***^(*B*)^, ***σ***^(*B*)^) are samplewise statistics. In general, 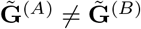. Due to samplewise standardization, the same raw vector can yield different normalized inputs across samples, acting as implicit data augmentation and inducing sample-dependent posteriors. This componentwise transformation preserves decomposability.

### 2.5 Gene-level probabilistic contributions

As mentioned above, the joint conditional distribution of a sample equals the joint conditional distribution of all gene vectors included in the sample:

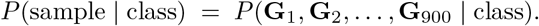

This is computed by the product node connecting each conditional distribution:

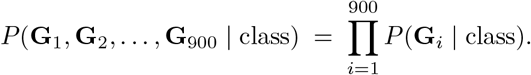

Therefore, at inference, we can extract any gene-level conditional distribution,

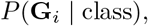

which represents the probabilistic contribution of that gene in the sample when all other genes are unobserved. For instance, in this paper we set cancer-cell and normal-cell classes; after optimization, the class-conditional distributions from a gene can be extracted as

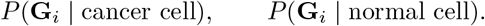

Here, *P* (**G**_*i*_ | cancer cell) represents the probabilistic contribution corresponding to the model’s decision process indicating the extent to which the gene supports classifying the sample as cancer and *P* (**G**_*i*_ normal cell) is interpreted analogously. If the model is successfully optimized without overfitting or underfitting on the training domain, it is expected that these class-conditional distributions reflect functional contributions to the specific class beyond the statistical frequency of the gene on the training domain.

### 2.6 Comparison of probabilistic contributions between genes

In our experiment, the probabilistic circuit is implemented in log-space. The log joint class-conditional density is

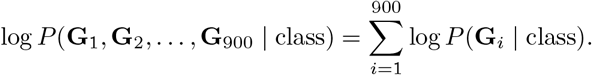

Here, we define the probabilistic contribution for a specific class as

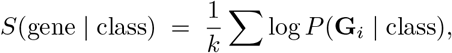

where *k* is the number of appearances of the gene across the extracted samples. In this paper, we define the cancer-cell and normal-cell contribution scores as

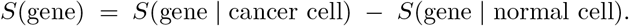

A high *S*(gene) indicates that the gene contributes more to the cancer class, whereas a low *S*(gene) indicates a stronger contribution to the normal class. By comparing this score across genes, we compare their probabilistic contributions.

### 2.7 Sample-level gene probabilistic contributions

Because of the sample-level standardization of the input vectors, the probabilistic circuit is trained to encode cancer-cell and normal-cell contribution scores for each gene under the genetic context of the sample. We therefore define the sample-level scores as

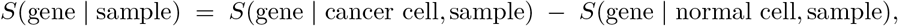

With

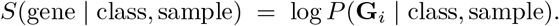

This sample-level gene score captures the sample dependency of gene function: for instance, genes with high *S*(gene | sample) in some samples but low *S*(gene | sample) in others suggest that the model considers those genes important for cancer classification only in specific gene contexts.

### 2.8 Datasets

We used the GSE131907 single-cell lung cancer atlas [18] including RNA-seq data from 208,506 cells spanning normal tissues and cancer from early to metastatic stages in 44 patients. Single-cell RNA sequencing was performed on whole cells derived from primary sites (tLung and tL/B), pleural effusions (PE), lymph nodes (mLN), and brain metastases (mBrain), as well as normal tissues from lung (nLung) and lymph nodes (nLN).

Two class labels—cancer cell and normal cell—are assigned to cells from tLung and nLung epithelium cells, respectively. The training, validation, and test sets were created by randomly splitting each subset into 3,420/570/570 cells per label at a fixed random seed. As described in Section 2.3, we randomly extracted 900 expressed genes from each cell at a fixed random seed; for each RNA, the corresponding transcription start site (TSS) was determined using the reference genome annotation in Gene Transfer Format (GTF). The selection was performed regardless of absolute RNA expression level.

In addition, to assess the robustness of the model across patients, we prepared additional training, validation, and test sets comprising data from different patients without any patient overlap. The numbers of cells in the training, validation, and test sets are 2,700, 500, and 1,000, respectively, including data from 7, 1, and 2 independent patients. We adopted Nucleotide Transformer v2 from InstaDeep and NVIDIA [8], which has 500 million parameters, and converted nucleotide sequences from −2000 bp upstream to +500 bp downstream of the TSS into 1,024-dimensional vectors. As a result, each cell has 900 gene vectors, and each gene vector represents a specific RNA in the sample.

### 2.9 Experimental Setting

We set the hyperparameters of our circuits to 2, 16, 2, and 2 for depth, latents, repetitions, and pieces, respectively, for the patient-mixed model, and 3, 24, 2, and 2 for the patient-independent model [12]. We trained the model with a batch size of 20 using the Adam optimizer[19].

### 2.10 Statistical Filtering

A key concern is overfitting to the statistical frequencies of gene types. To address this, we filtered genes based on the following criteria:

#### Contradictory count–score pairs

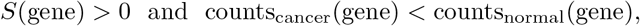

or

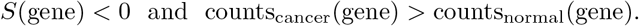

Such filtering reduces the influence of raw frequency in the dataset on the learned contributions.

## 3 Results and Discussions

### 3.1 Assessment of feature representations during optimization

As mentioned in the method section, DVPNet is designed so that its decision process or the class-conditional probabilistic distribution for each gene vector can be visualized. However, this does not guarantee that the model focuses on and captures meaningful feature representations for biological analysis. First, similar to most deep-learning-based classification models, the model needs to avoid the following situations:

- **Underfitting**. Here, the training task is too complex for the model to capture meaningful features for the classification task due to insufficient model capacity, inappropriate representations, or noisy supervision. In this situation, the model cannot achieve sufficient classification performance on the training dataset.
- **Overfitting**. Here, the model does not show robustness. The model can show sufficient performance only on the training dataset but not on the test dataset. This suggests that the model captures overly complex feature representations meaningful only for the training dataset. For meaningful biological analysis, at least the model needs to grasp generalized features applicable to the target domain (e.g., a test dataset split from the entire dataset).

The overall classification performance of the model on the training and test datasets is as follows. The patient-mixed model achieved AUROC, AUPRC, maximum accuracy, and F1 scores of 0.992, 0.994, 0.974, and 0.974 on the training set, and 0.975, 0.981, 0.940, and 0.939 on the test set, respectively. For the patient-independent model, it achieved AUROC, AUPRC, maximum accuracy, and F1 scores of 0.999, 0.999, 0.991, and 0.991 on the training set, and 0.976, 0.979, 0.920, and 0.918 on the test set.

The high performance on the training set indicates that the model does not exhibit underfitting during optimization. In addition, the relatively small performance gap between the training and test sets suggests that the model does not suffer from severe overfitting and generalizes well to unseen combinations of genes.

While the model avoids underfitting and overfitting, this is not sufficient to guarantee the advantage of our model for biological analysis compared to other statistical methods. The classification task should be solved using the feature representations of the embedding vectors from the Nucleotide Transformer as well as label-wise gene occurrence frequencies.

In Figure 2, the statistical frequency counts between two labels on the training set are compared with the mean *S*(gene) score provided by the model. The moderate Pearson (*r* = 0.356) and Spearman (*ρ* = 0.408) correlations between gene occurrence frequency differences and probabilistic contribution scores suggest that gene frequency alone does not fully explain the learned contributions. This observation is consistent with the model leveraging additional information encoded by the Nucleotide Transformer embeddings during classification.

**Figure 2.**
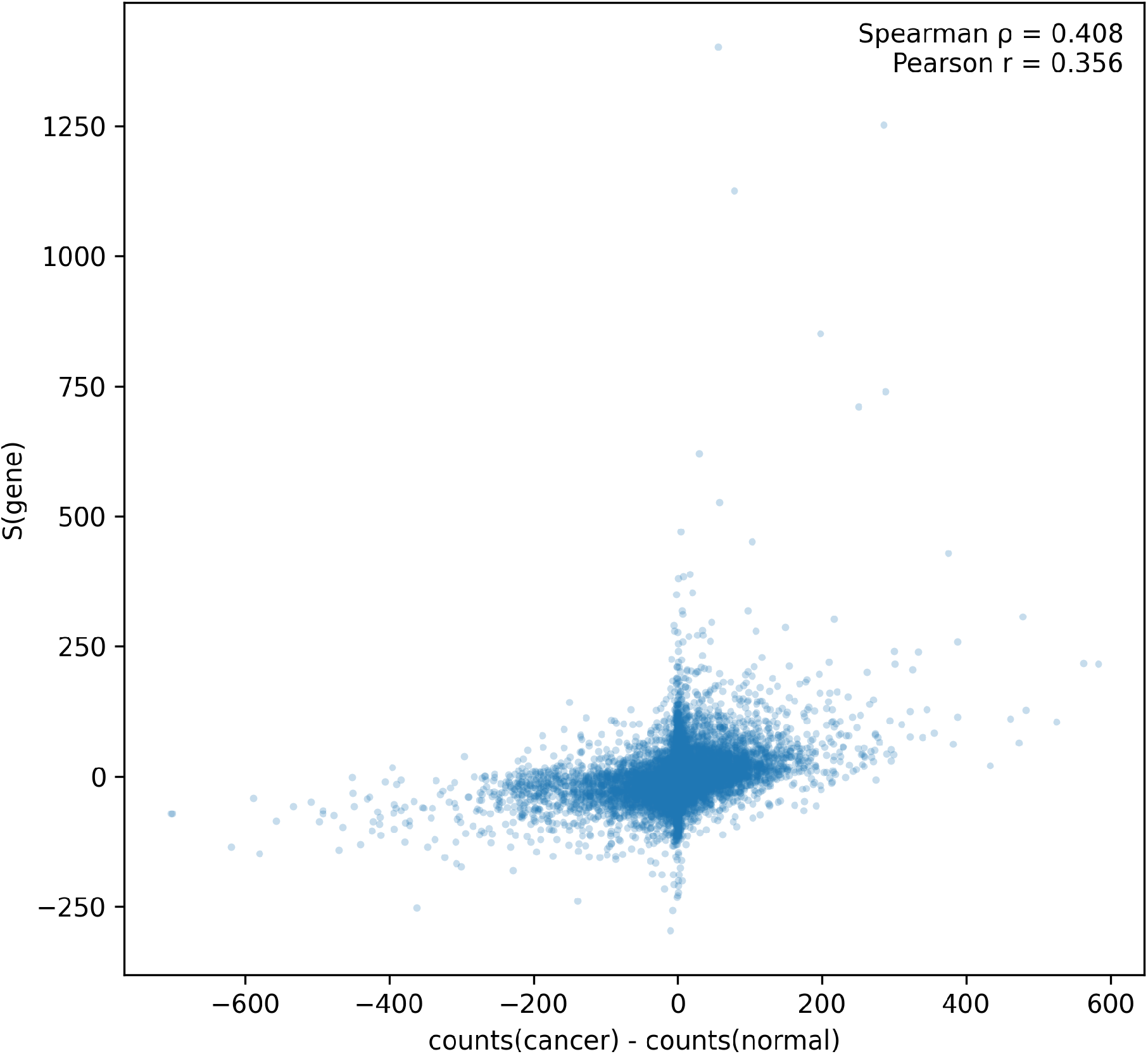
Relationship between label-wise gene occurrence count differences and the mean probabilistic contribution score *S*(gene). Pearson’s *r* and Spearman’s *ρ* are reported in the plot.

In summary, the experimental results suggest that the model was successfully optimized without suffering from underfitting or overfitting, and the probabilistic contributions of genes were determined based on a combination of the statistical frequency and the feature representations from the Nucleotide Transformer. Assuming that the Nucleotide Transformer encodes inherent biological functions, these probabilistic contributions have the potential to enable meaningful biological analysis and provide new biological insights.

### 3.2 A WGCNA-based network analysis

Weighted gene co-expression network analysis (WGCNA) [20] [21]is a widely used method to identify clusters of genes whose RNA expression levels are highly correlated. This analysis contributes to understanding and exploring the system-level functionality of genes [21]. Normally, WGCNA is constructed based on correlations in RNA expression variation across samples; however, instead of using RNA expression levels, we utilized the sample-dependent probabilistic contribution, *S*(gene | sample), to construct a new genetic network. Unlike traditional WGCNA, this network depends on the model’s decisions regarding whether a given gene positively or negatively contributes to *S*(gene) for a specific sample.

Using all training samples and their corresponding *S*(gene | sample) values for 9,540 genes, an unsigned weighted network was constructed with a soft-thresholding power automatically selected according to the scale-free topology criterion. Gene modules were identified from the topological overlap matrix using hierarchical clustering followed by dynamic tree cutting (minimum module size = 2). This procedure identified 50 distinct modules comprising 1,812 genes, while the remaining genes were excluded as noise. Within each module, genes were ranked by intramodular connectivity, defined as the sum of connection strengths to other genes in the same module, and genes with the highest intramodular connectivity were considered hub genes.

To characterize the biological functions represented by each identified module, Gene Ontology (GO) enrichment analysis [22] [23] was performed separately for each module using the genes assigned to that module. Enriched GO terms were used to interpret the functional roles of individual modules and their corresponding hub genes.

Table 1 summarizes the module-level scores and representative hub genes, which are determined based on intramodular connectivity. Modules comprise between 11 and 301 genes and are ordered by *S*(genes) computed across all samples, representing the overall probabilistic contribution to the cancer cell class scored by the classification model. The module with the highest *S*(genes) value (45.71) is the orange module, whereas the royal blue module shows the lowest *S*(genes) value (−43.90). To illustrate condition-specific probabilistic contributions based on class labels, *S*(genes) values were computed separately for samples with cancer cells as ground truth and for samples with normal cells as ground truth. The 0074ff module shows the largest difference between *S*(genes | cancer samples) and *S*(genes | normal samples) (38.62), suggesting that the model assigns a stronger contribution to this module for cancer classification in cancer samples, and conversely for normal samples.

**Table 1:**
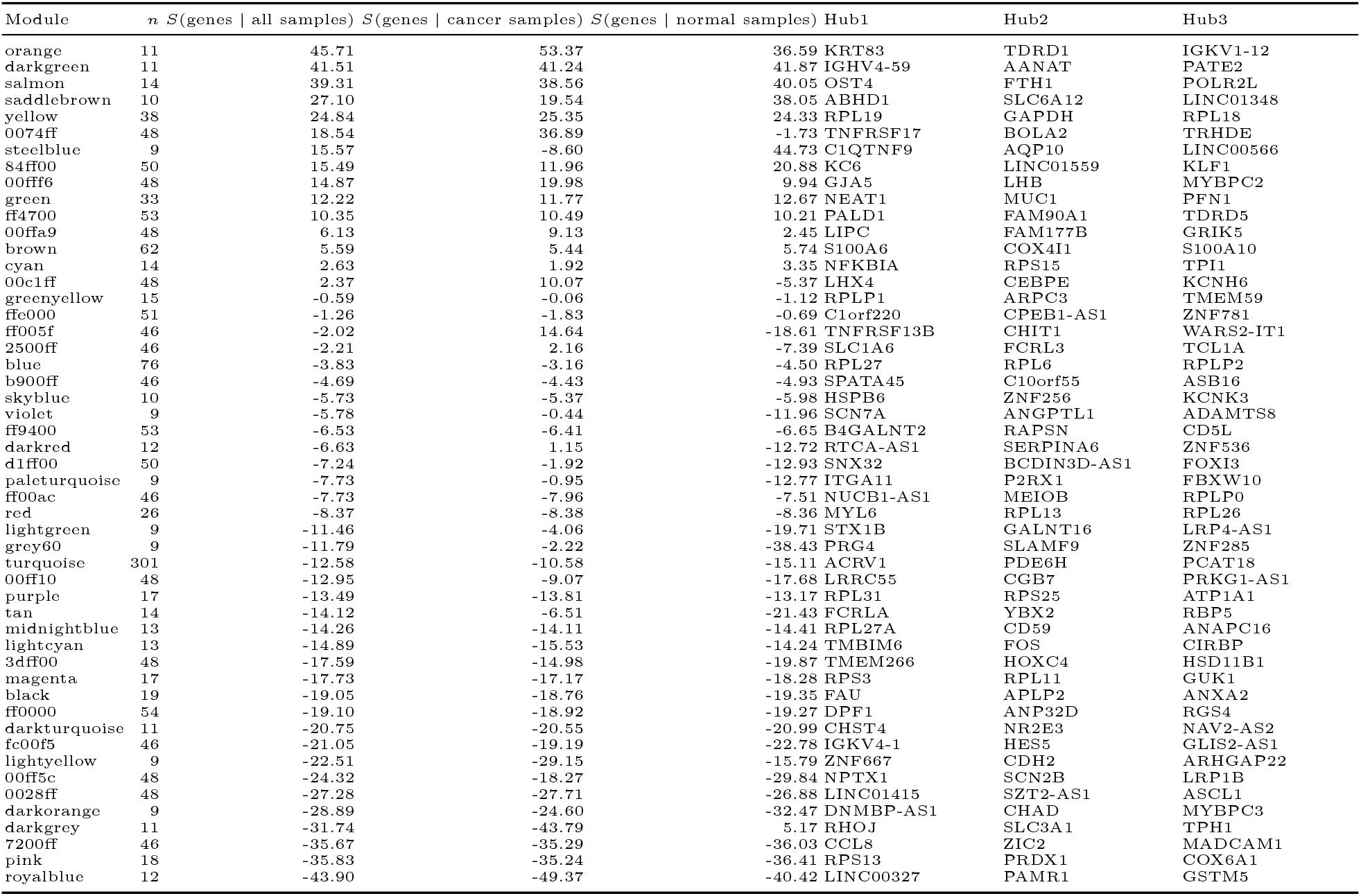
Module-level scores and hub genes. Modules are ordered by *S*(genes | all samples) (descending).

Table 2 presents the five representative Gene Ontology enrichment terms for each module. These terms summarize the dominant biological functions associated with each module.

**Table 2:**
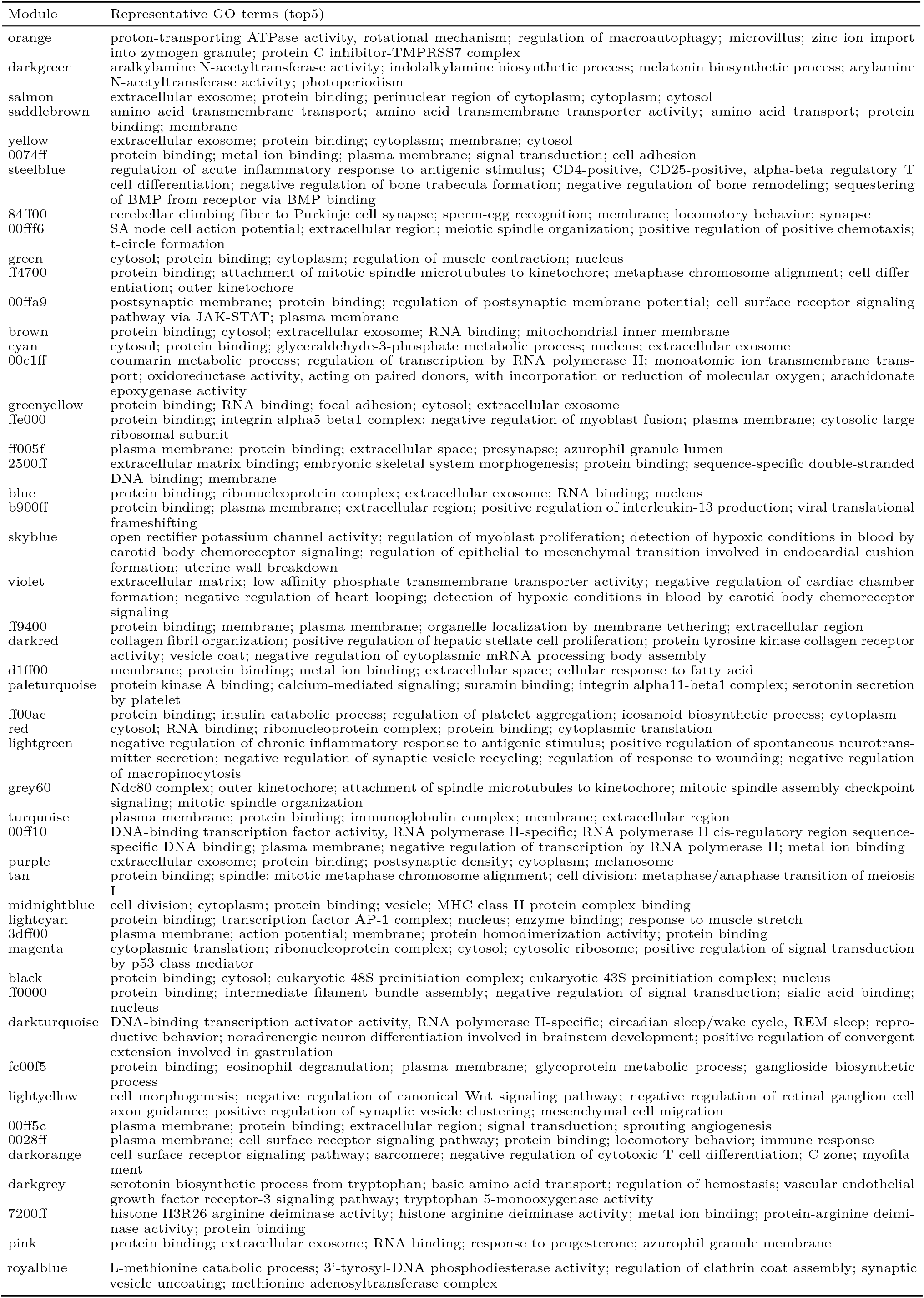
Representative GO enrichment terms (top5) for each module (ordered by *S*(genes | all samples) descending).

This explainable AI based network may provide biological insights distinct from those obtained by traditional WGCNA, which is constructed solely from correlation relationships of RNA expression levels. Because the proposed network is not derived directly from RNA expression correlations, the resulting *S*(genes) scores reflect probabilistic contributions inferred from the model’s decision process. Consequently, the genetic modules and their associated biological pathways, as characterized by Gene Ontology analysis, may highlight functionally relevant gene sets that contribute to distinguishing cancer cells from normal cells.

### 3.3 Analysis of pathway level contribution

In this section, we applied GO enrichment analysis to 9540 genes on the training set without any clustering analysis. Then, the derived biological terms were evaluated based on mean S(genes) score averaged from S(genes) of all the genes comprising the pathway. This analysis can provide information about which biological pathways are more contributed to the distinguish cancer cells from normal cells for the classification model.

GO enrichment analysis identified a total of 4,000 biological terms with q-values less than 0.05. GO terms were ranked according to their mean S score (*S*_mean_) calculated from the overlap genes.

The top 100 GO terms with the highest *S*_mean_ values are listed in Table 3, and the corresponding overlap genes for each term are shown in Table 4. These top-ranked GO terms include multiple entries related to immunoglobulin complexes, complement activation, antibody-dependent cellular cytotoxicity, and humoral immune response pathways. The bottom 100 GO terms with the lowest *S*_mean_ values are presented in Table 5, and their corresponding overlap genes are summarized in Table 6. These GO terms span a wide range of functional categories, including immune regulation, complement-related processes, metabolic pathways, ion transport, and cellular structural components. All of these terms exhibit negative *S*_mean_ values relative to other enriched GO terms.

**Table 3:**
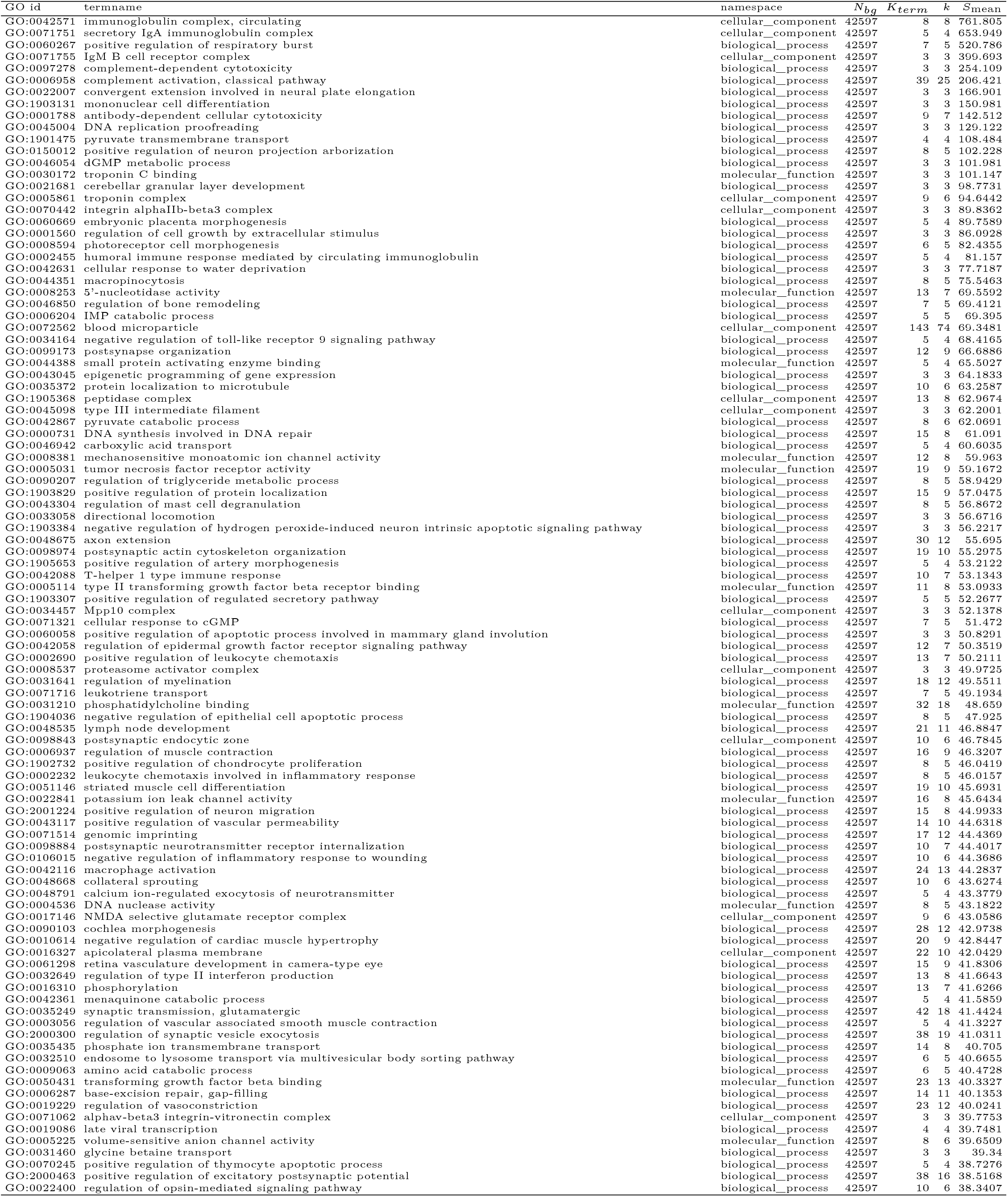
Top GO terms by S(mean)

**Table 4:**
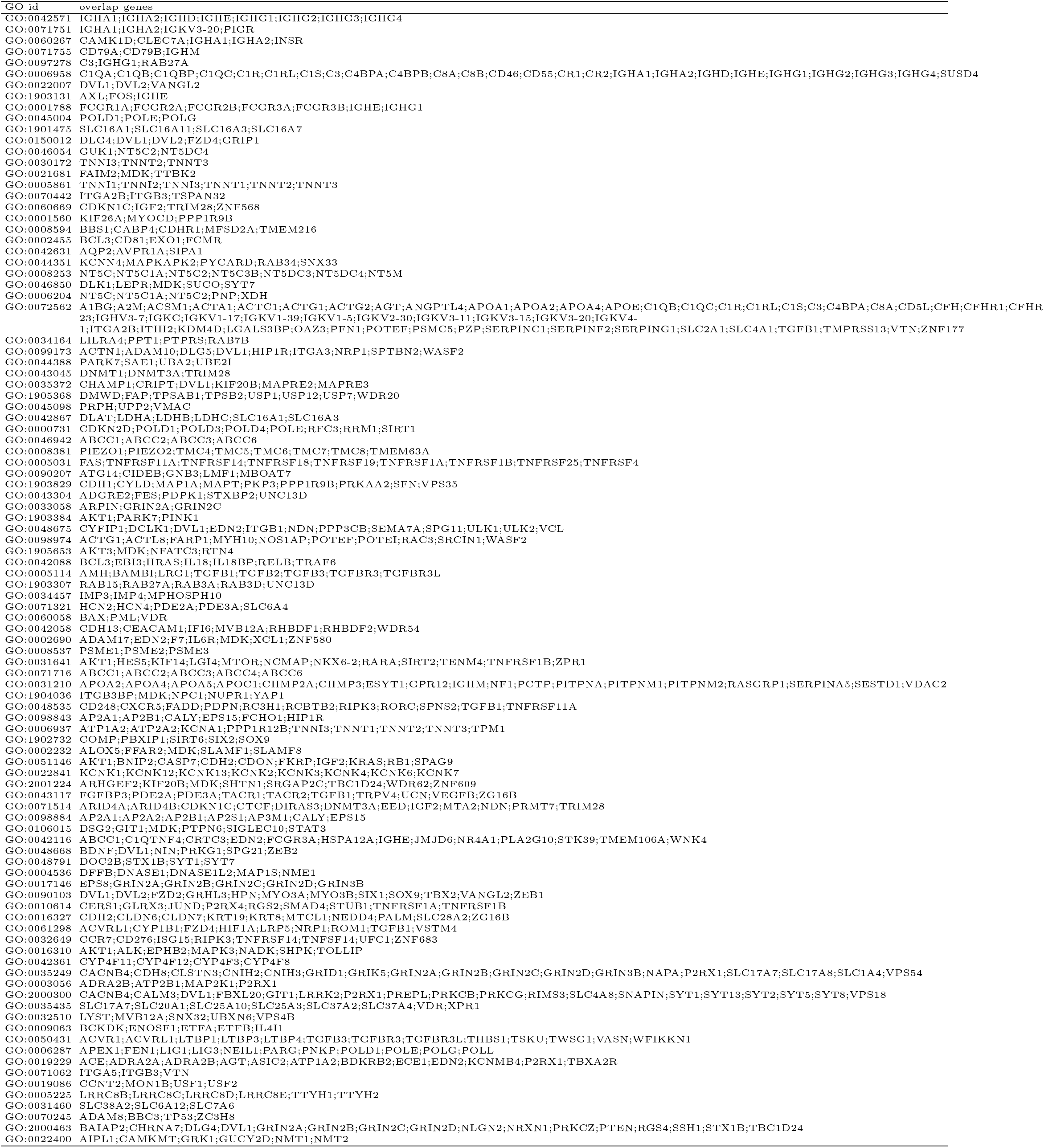
Overlap genes for top GO terms.

**Table 5:**
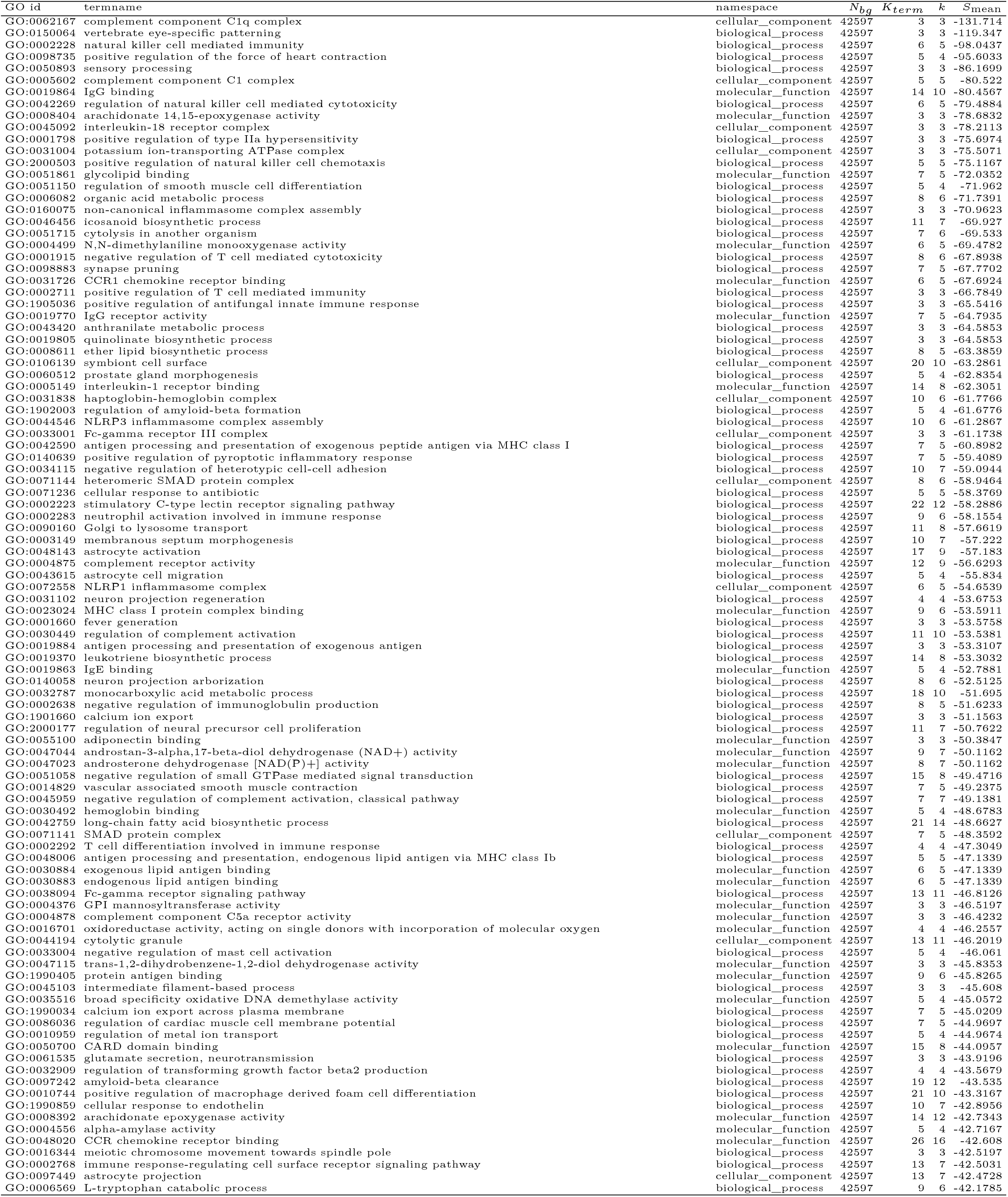
Bottom GO terms by S(mean)

**Table 6:**
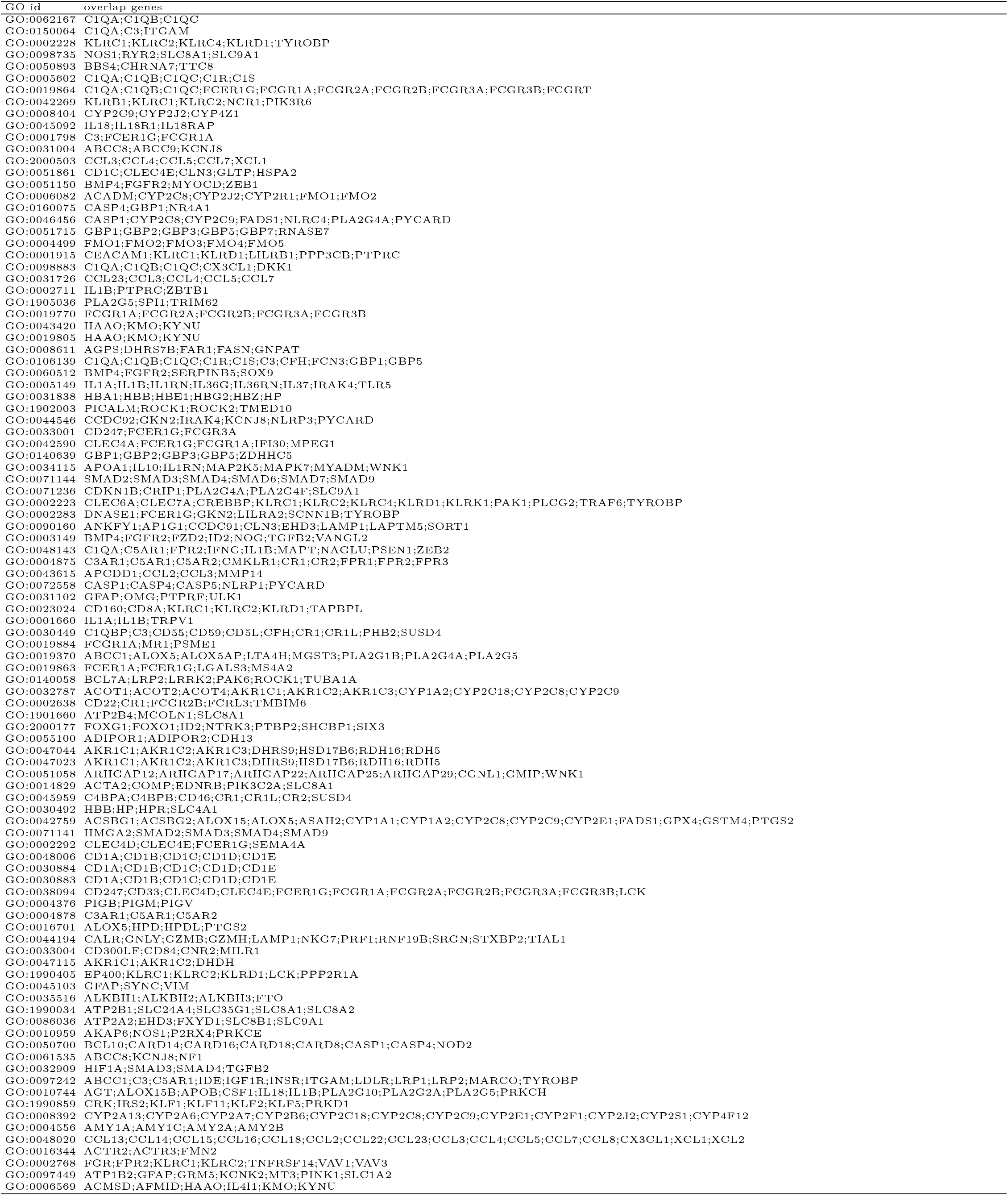
Overlap genes for bottom GO terms.

### 3.4 Analysis beyond statistics

In this section, we analyzed 1,524 genes from the training set that exhibited contradictory count–score pairs (Section 2.10), namely genes with *S*(gene) >0 while counts_cancer_ <counts_normal_, or *S*(gene) <0 while counts_cancer_ >counts_normal_. Table 7 and Table 8 display the top 100 and the bottom 100 genes based on *S*(gene) respectively.

**Table 7:**
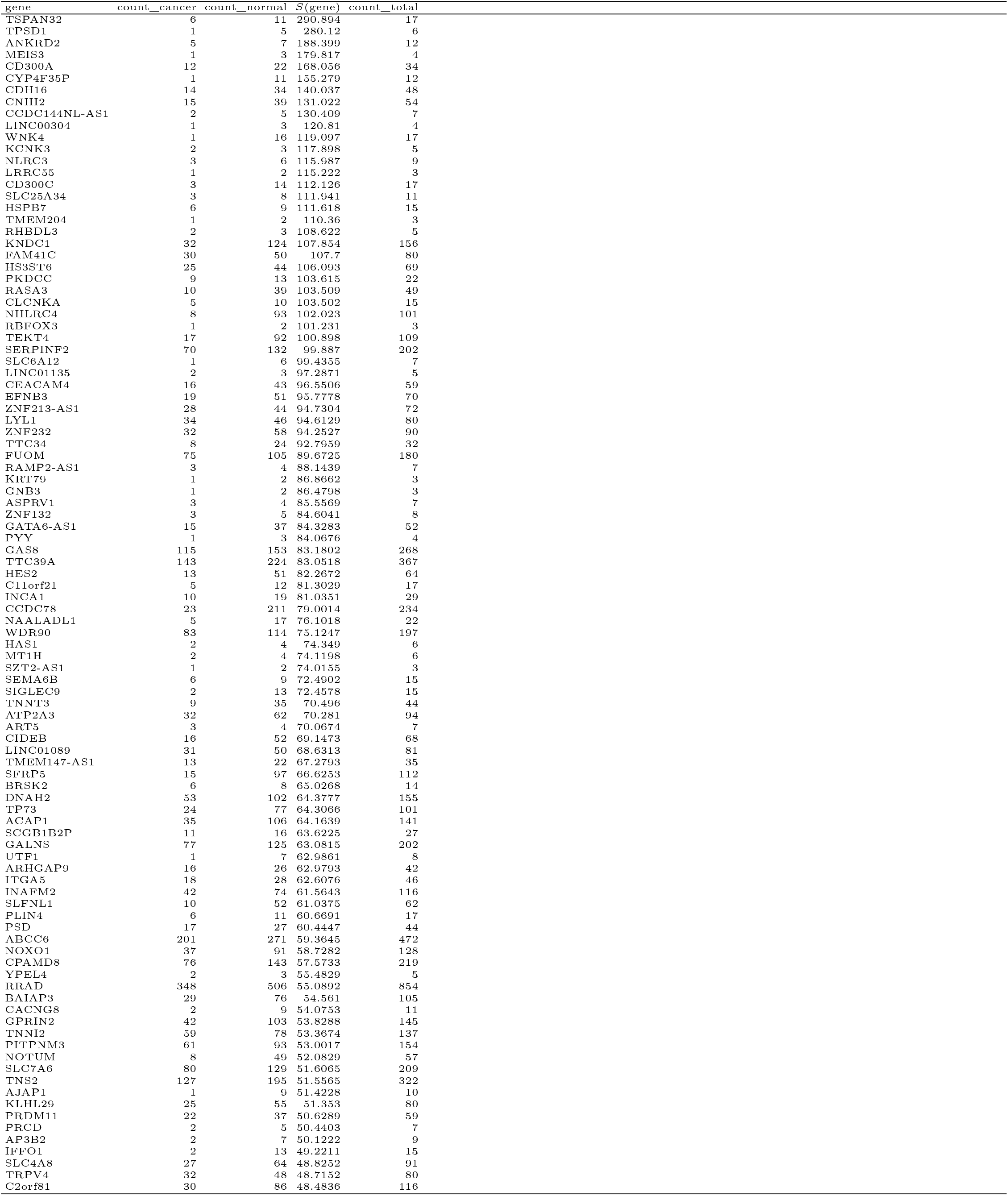
Top 100 genes among the 1,524 genes exhibiting contradictory count–score pairs.

**Table 8:**
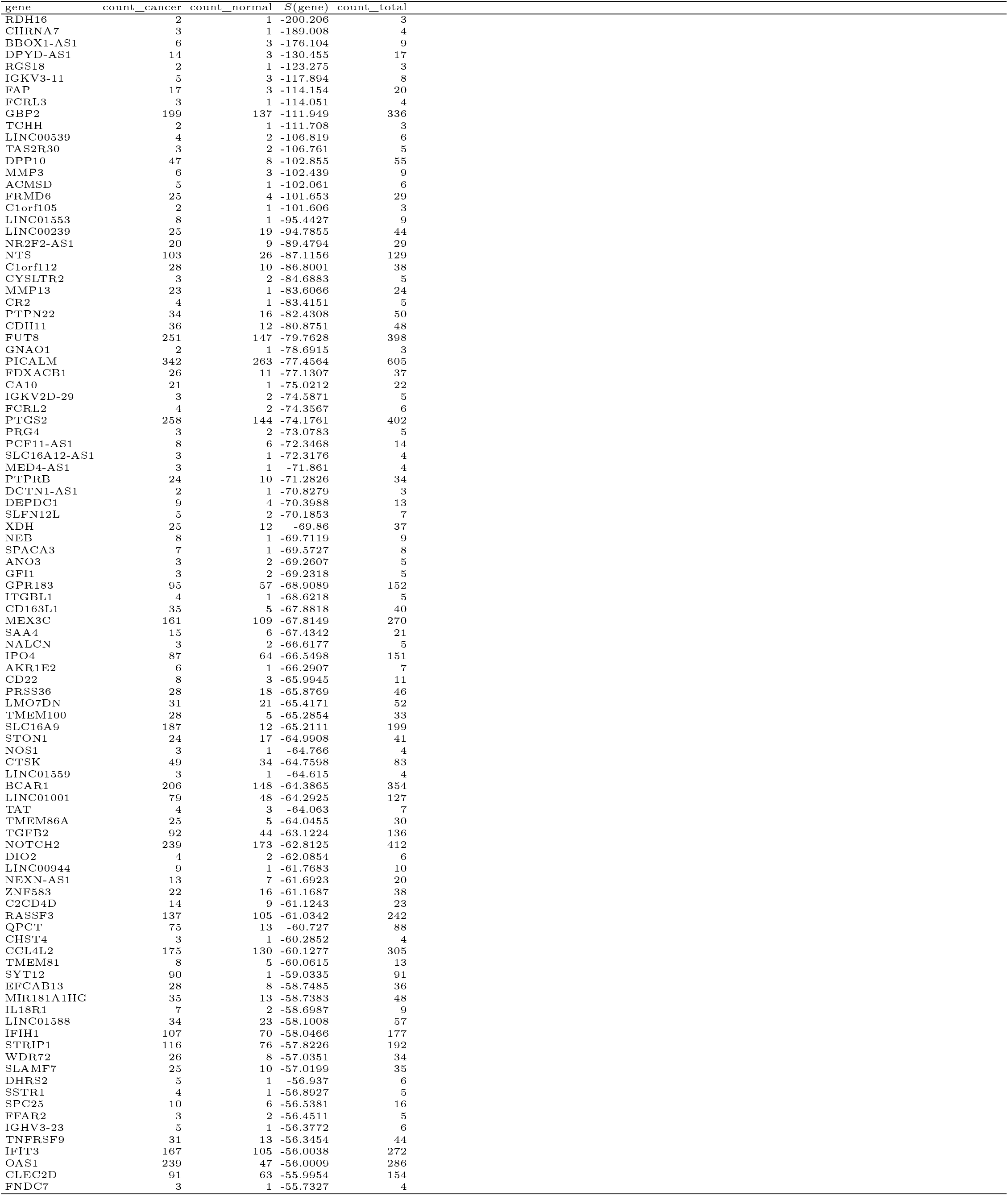
Bottom 100 genes among the 1,524 genes exhibiting contradictory count–score pairs. $.

The classification model is provided with only two sources of information: gene occurrence statistics in the training set and feature representations encoded by the Nucleotide Transformer. If the model relied solely on statistical gene counts, genes with higher occurrence in a given class would naturally receive higher contribution scores for that class. However, the existence of these contradictory genes demonstrates that this is not the case.

Because the statistical signal alone cannot explain the observed score assignments for these genes, the results indicate that the feature representations encoded by the Nucleotide Transformer dominate and override the frequency-based signal for the corresponding class decisions. Focusing on such genes enables further biological analysis that is not directly driven by gene occurrence statistics.

It is natural to consider that expressed genes appearing more frequently in cancer cell samples functionally contribute more to cancer cells. However, this result suggests that the explainable AI model evaluates the differentiation between cancer cells and normal cells as being regulated by more complex relationships beyond simple statistics; that is, the importance of genes for cancer cells cannot be determined solely by their occurrence statistics.

This result supports that our model can comprehensively evaluate probabilistic contributions by integrating both statistical information and feature representations encoding biological functions. As discussed above, this enables analyses beyond simple gene occurrence statistics and provides a complementary perspective to traditional statistics-based approaches, such as WGCNA and differential gene analysis [24].

### 3.5 Biological validation of the genetic profiling

Wet-lab experiments to test whether the genes prioritized by our probabilistic contribution scores exhibit functional relevance in cancer biology. If such experiments validate that top-ranked genes (e.g., TSPAN32, TPSD1, ANKRD2) are functionally involved in tumor progression or immune modulation, this would strongly support the biological relevance of our framework. However, such experimental validation was beyond the scope of the present study. Instead, we examined existing literature to assess whether previously reported clinical or biological findings are consistent with the genes prioritized by our framework.

Among the top-ranked genes in Table 7, multiple genes have been independently investigated as therapeutic targets in cancer research, including ITGA5[25][26], SIGLEC9[27][28], and NOTUM[29][30]. In addition, several well-characterized genes involved in tumor suppression and stress-response pathways, such as TP73[31], SFRP5[32], and NLRC3[33], are also present among the top-ranked genes. Furthermore, key regulators of immune and microenvironmental signaling including CD300A/C[34][35], PITPNM3[36], and SEMA6B[37] are included in the list.

The consistency between previously reported findings and our probabilistic gene ranking suggests that the proposed framework successfully prioritizes genes with established relevance in cancer biology. Importantly, the presence of less-characterized genes within the same ranked list indicates that our framework may also highlight novel candidates warranting further investigation.

### 3.6 Validation of Random Gene Selection

In this paper, we randomly selected 900 expressed genes for each cell sample and converted each gene into an embedded vector using the Nucleotide Transformer, while treating all genes equally regardless of their RNA expression levels. This assumes that genes with lower RNA expression levels may also contribute importantly to biological regulation or functionality. As discussed in the previous sections, our model avoids both overfitting and underfitting and shows that the probabilistic contribution scores are determined beyond simple statistics, supporting the validity of our approach.

However, this approach is not the only possible way to use DVPNet. Other approaches may provide different biological insights. For instance, we can select the top *k* genes for each sample based on RNA expression levels, or incorporate RNA expression information into the model by mathematically modifying the embedding vectors according to the expression levels, without affecting the smoothness and decomposability of the probabilistic circuits. Under these alternative approaches, the model would be optimized using different sources of information. Such optimization may assign probabilistic scores to genes from different biological perspectives by leveraging RNA expression level information. Exploring more effective approaches remains a task for future research.

### 3.7 Limitations

In this study, we used the GSE131907 single-cell lung cancer atlas for the explainable model to extract meaningful representations to differentiate between cancer cells and normal cells. However, this setting may involve some overestimation. First, we utilized only epithelial cell samples from primary sites labeled tLung and nLung to avoid situations in which the two classes have different cellular origins. For instance, if samples in the cancer-cell class are derived from epithelial cells, while samples in the normal-cell class are derived from lymph nodes, the model may focus on differences between epithelial cells and lymph cells rather than differences between cancer cells and normal cells. Our selection avoids this situation, but it also introduces a limitation.

Samples labeled as the cancer-cell class may include cells derived from the tumor microenvironment in addition to cancer cells. In this case, the classification task between cancer cells and normal cells may be implicitly shifted to a classification task between the tumor microenvironment and the normal tissue microenvironment. This interpretation is supported by our results in Section 3.3, where the top-ranked GO terms represent immune-related functions, including immunoglobulin complexes, complement activation, antibody-dependent cellular cytotoxicity, and humoral immune response pathways, suggesting that the model captured differences in immune responses between the tumor microenvironment and the normal tissue microenvironment.

Second, as our training dataset includes only one cancer type (lung cancer), drawing conclusions about biological differences between general cancer cells and normal cells would require a broader dataset encompassing multiple cancer types and corresponding normal cells.

## 4 Conclusion

In this paper, we presented a new XAI-based genetic profiling approach in which an explainable classification model assigns a probabilistic contribution score to each gene based on optimization for the classification task between cancer cells and normal cells. The model achieved high accuracy on both the training and test sets without exhibiting underfitting or overfitting, suggesting that the proposed classification framework is valid and that the model was successfully optimized. In addition, this study shows that the model utilized both statistical signals and biological feature representations encoded by the Nucleotide Transformer. Some genes exhibited probabilistic contributions that were contradictory to their statistical frequencies, suggesting that our analysis, which integrates a biological foundation model with statistical information, may provide biological insights different from those obtained through traditional statistics-based analyses.

